# Sex differences in the relationship between social difficulties and executive dysfunction in children and adolescents with autism spectrum disorder

**DOI:** 10.1101/501932

**Authors:** Tonje Torske, Terje Nærland, Daniel S. Quintana, Ruth Elizabeth Hypher, Anett Kaale, Anne Lise Høyland, Sigrun Hope, Jarle Johannessen, Merete G Øie, Ole A Andreassen

## Abstract

The prevalence of autism spectrum disorder (ASD) in boys is nearly four times higher than in girls, and the causes of this sex difference are not fully known. Difficulties in executive function may be involved in development of autistic symptomatology. Here we investigated sex differences in the relationship between executive function in everyday life and social dysfunction symptoms in a sample of 116 children (25 girls) aged 5-19 years with IQ above 70 and with a diagnosis of ASD. They were assessed with the Behavior Rating Inventory of Executive Function (BRIEF) and the Autism Diagnostic Interview Revised (ADI-R). We found no significant differences in BRIEF or ADI-R scores between girls and boys after correcting for multiple testing. Nested linear regression models revealed significant sex differences in the relationship between executive function and both reciprocal social interaction (*p*<0.001) and communication (*p*=0.001) over and above the main effects of age, sex, IQ and comorbid attention deficit/hyperactivity disorder diagnosis. We did not find sex differences in the relationship between executive dysfunction and restricted and repetitive behaviors. Altogether, our results provide a greater understanding of the sex-specific characteristics of ASD and may suggest that boys and girls can benefit from different intervention strategies.

## Introduction

Autism spectrum disorder (ASD) is overrepresented in boys compared to girls. Traditionally, the male-to-female ratio is thought to be 4:1 ^1^. However, a recent meta-analysis of population based ASD studies concluded that the male-to-female ratio is closer to 3:1, suggesting that researchers and health professionals may currently overlook ASD in females ^2^. This sex difference influences the identification of autistic symptoms and obtaining an accurate diagnosis, as well as intervention options and the provision of suitable resources and services for people with ASD ^3^. Furthermore, the underlying causes of the difference in ASD occurrence between boys and girls are not fully known. Research on females with ASD has been limited, and most of the literature on ASD is based on boys and young men ^4^. Thus, there is a growing need for a better understanding of the sex differences in ASD and there is an increased research focus on girls with ASD ^1,3,5^.

Some studies suggest that the sex difference in ASD prevalence can partially be due to sex-differential genetic and hormonal factors ^6^. However, the genetic factors underlying the skewed sex ratio in ASD remains mostly unknown, and cannot be explained by X-linked variants since most known ASD risk genes are located in autosomal regions ^6^. There is some evidence for increased mutational burden in females and their families, which indicates an elevated threshold for developing ASD in girls ^7^. This has been interpreted as a female protective effect, in other words, a greater resistance to ASD from genetic causes in females ^8^. Even though there are no complete molecular explanations for this hypothesis ^9^, studies suggest that the male bias is most likely due to female protective factors rather than male-specific risk factors. A possible consequence of an increased genetic load in girls, is that those who reach a clinical diagnosis of ASD often have lower intelligence and more behavioral problems than boys with ASD ^10^.

An important factor in the prevalence ratio in ASD seems to be related to cognitive level; lower IQ is associated with a lower male-to-female ratio ^2,5,11^. However, this finding needs to be treated with caution, since only about half of the studies included in Loomes and colleague’s recent review included sufficient information regarding IQ ^2^. It has also been recognized that the autistic symptoms are less apparent in girls than boys. This phenomenon might be due better learning of compensatory behaviors and skills to mask their social challenges ^12,13^ and that parents, teachers, and clinicians are less able to recognize autistic symptoms in girls ^14^. Girls with ASD tend to have better social skills and less behaviour problems than boys with ASD, which might make it harder to recognise their autistic characteristics ^10^. Furthermore, some have found that girls with ASD have less repetitive behavior and interests compared to boys with ASD ^1,15^.

EF deficits constitute one of the main cognitive theories of ASD ^16–18^, together with Theory of Mind and Central Coherence ^19^. Recent meta-analyses confirm that on average, people with ASD perform worse on executive function (EF) tasks than neurotypical controls ^20,21^. EF comprises several components including inhibition, working memory, flexibility, emotional control, initiation, planning, organization, monitoring and self-control ^17,22^. These components enable the individual to disengage from the present context to effectuate future goals. Demetriou and colleagues ^20^ found consistent evidence of an overall moderate effect size (Hedges’ *g* = 0.48) of executive dysfunction in ASD, that the deficits are relatively stable across development, with few differences across subdomains ^20^. In a meta-analysis that also included children and adolescents with ASD and comorbid attention deficit/hyperactivity disorder (ADHD), Lai and colleagues ^21^ confirmed that children with ASD tend to have executive dysfunction with small to moderate effect sizes (Hedges’ *g* = 0.41-0.67), and that this was not solely accounted for by the effect of comorbid ADHD or general cognitive abilities. Further, the questionnaire Behavior Rating Inventory of Executive Function (BRIEF) was found to be a better clinical marker of ASD than performance based tests ^20^. This is probably because it can be difficult to generalize from EF assessed in highly structured laboratory settings, and that questionnaires regarding everyday functioning have a higher ecological validity and thus also a better clinical utility than neuropsychological tests ^20,23^. In addition, intelligence and age are factors that might influence EF in children with ASD ^24^.

Sex differences in the relationship between EF and social function might contribute to the skewed sex distribution in ASD. If girls who reach a clinical diagnosis of ASD tend to be more impaired and have a higher genetic burden than boys ^10^, the relationship between EF deficits and social difficulties may also be different in girls. Studies investigating this relationship have focused on specific subdomains of EF examined mainly by neuropsychological tests ^11,25–27^. Since some EF difficulties may not be observable in a laboratory setting, informant based measures and questionnaires like the BRIEF might add valuable information ^23^. In addition to EF, there are indications that there also may be sex differences in people with ASD within domains such as mentalizing, emotion perception, perceptual attention to detail, and motor function ^4^.

Although studies have identified a relationship between key ASD traits, such as social dysfunction, and EF ^28^, there are few studies focusing on how *sex* might impact the relationship, and the findings have been inconsistent. Some studies have indicated that females with ASD have more impairment in EF compared to males ^25^. In a relatively small group of participants, Lemon and colleagues ^25^ found that only girls showed poorer response inhibition. Others have reported that females with ASD outperform males on executive tasks related to processing speed and verbal fluency ^11,26^.

With regard to everyday functioning in children with ASD, there is one study, to our knowledge, of sex differences in the relationship between the Autism Diagnostic Interview Revised (ADI-R) and adaptive behavior ^29^, and another study of sex differences in parent-reported EF and adaptive behavior ^30^. Mandic-Maravik and colleagues ^29^ found different associations of autistic symptoms with various aspects of adaptive behavior between the sexes. White and colleagues ^30^ reported a correlation between EF difficulties and decreased adaptive ability in both males and females. However, females had more EF difficulties on the BRIEF and more difficulties on the Daily Living Skills domain on the Vineland Adaptive Behavior Scales. To the best of our knowledge there are no studies of how sex differences influence the relationship between parent-rated EF in everyday life (BRIEF) and autistic symptomatology (ADI-R).

The main aim of the current study was to investigate the relationship between EF in everyday life rated by parents and autistic symptomology, and to investigate possible sex differences in this relationship. In accordance with the female protective effect hypothesis, that girls would need to be more impaired to have the same amount of ASD symptoms as boys, we hypothesized the relationship between EF deficits and autistic symptomology to be stronger in girls than boys.

## Methods

### Participants

The participants were recruited from Norwegian health services specializing in the assessment of ASD and other neurodevelopmental disorders. The study was part of the national BUPgen network ^31^. The current sample consisted of 25 girls and 91 boys with ASD who were recruited between 2013 and May 2018 and assessed at age 5-19 years. Fifteen of the children (2 girls, 13 boys) were diagnosed with childhood autism, 9 (2 girls, 7 boys) with atypical autism, 57 (14 girls, 43 boys) with Asperger syndrome and 35 (7 girls, 28 boys) with unspecified pervasive developmental disorder (PDD-NOS).

The male:female ratio was 3.6:1. In total, 40 children (34.5%) had a comorbid disorder of ADHD. All participants had an intelligence quotient (IQ) within the normal range based on a standardized Wechsler’s test (Full-scale IQ ≥ 70) and spoke Norwegian fluently. Exclusion criteria were significant sensory losses (vision and/or hearing).

### Clinical assessment

The children were assessed by a team of experienced clinicians (clinical psychologists and/or child psychiatrists and educational therapists). Diagnostic conclusions were best-estimate clinical diagnoses derived from tests, interview results and observations. All diagnoses were based on the International Statistical Classification of Diseases and Related Health Problems 10^th^ Revision (ICD-10) ^32^ criteria, and the autistic symptoms were evaluated using the Autism Diagnostic Observation Schedule (ADOS) ^33^ and/or Autism Diagnostic Interview-Revised (ADI-R) ^34^. In addition, the assessment included a full medical and developmental history, physical examination and IQ assessment. Because ASD and ADHD often co-occur (29), the current study also included children with ASD and comorbid ADHD.

For a subsample of the group n = 34 (10 girls), we also had neuropsychological test data from the Delis-Kaplan Executive Function System (D-KEFS) ^35^. We used five of the subtests from D-KEFS. Results are reported as mean scaled scores and standard deviations (10+/-3): 8.48 (3.44) for Trail Making Test Condition 4 Number-Letter Switching (n = 31), 8.97 (2.42) for Verbal Fluency Letter Fluency (FAS) (n = 32), 8.59 (3.39) for the Color-Word Inhibition Time (n=34), 8.44 (3.31) for Color-Word Inhibition/ Switching Time (n = 32), 10.39 (3.59) for Twenty Questions Initial Abstract Score (n = 28) and 10.19 (2.34) for Tower Test Total Achievement Score (n = 31). The subsample with neuropsychological test results was on average older than the total sample (11.8 versus 10.3 year; *p* = 0.009), and fewer had comorbid ADHD than the total sample (*p* = 0.043). However they did not differ from the total sample in sex distribution (*p* = 0.185). Due to a small sample size, the neuropsychological test results are included to describe the group and were not used in further analyses.

### Measures

#### Autistic symptoms

Autism Diagnostic Interview-Revised (ADI-R) diagnostic algorithm was used to assess autistic symptoms. The ADI-R is a clinical diagnostic tool based on a comprehensive interview with parents or primary caregivers of the child/ adolescent ^36^. The interview consists of 93 questions, and a predetermined number of these scores go into a diagnostic algorithm. The interview and scoring follow standardized procedures, and the interviewer records and codes the informant’s responses. The algorithm is divided into three functional domains based on the diagnostic criteria (qualitative deviations in): A = Reciprocal Social Interaction, B = Communication, C = Restricted, Repetitive, and Stereotyped Behavior. Higher scores indicate that an individual has a greater number of items representing core ASD deficits and/or more severe symptoms ^37^. All the participants were verbal children, and therefore the algorithm for verbal children was used. We used the Norwegian translation of the ADI-R ^38^.

#### Executive function (EF)

In order to assess EF parents completed the parent version of the BRIEF ^39^. The BRIEF for children and adolescents aged 5 to 18 years includes 86-item parent and teacher forms that allow professionals to assess everyday EF in the home and school environments ^39^. The BRIEF contains eight scales that are grouped in a Behavioral Regulation Index (BRI): Inhibit, Shift and Emotional Control, and a Metacognition Index (MI): Initiate, Working Memory, Plan/Organize, Organization of Materials and Monitor. *T-scores* of ≥ 65 are considered to represent clinically significant areas. The Global Executive Composite (GEC) is a summary score that incorporates all eight clinical scales. The GEC has high reliability in both standardized and clinical samples (Cronbach’s alpha = 0.80-0.98). The current study used the Norwegian version of the parent rating form, which has been reported to have high internal consistency (Cronbach’s alpha = 0.76-0.92) ^40^. Similar levels are described for the English version (Cronbach’s alpha = 0.80-0.98) ^39^.

#### Intelligence Quotient (IQ)

IQ was assessed using age-appropriate full-scale Wechsler tests of intelligence ^41–43^. We used the Norwegian versions of the Wechsler tests, which have Norwegian and/or Scandinavian norms ^44–46^.

### Statistical analyses

Analyses were conducted using the R statistical environment (version 3.5.0) using the “jmv” (Version 0.7.3.1; ^47^) and “cocor” packages ^48^. Statistical significance was set at *p* < 0.05 and adjusted according to number of comparisons. When adjusting critical p-values for multiple tests it is important to carefully consider the risks of type-I and type-II errors ^49^. Thus, we provide justifications below for how we adjusted tests for multiple comparisons to control the Type-I error rate. Conventional values were used for interpreting effect sizes (Effect size values of 0.2, 0.5, and 0.8, were considered small, medium, and large effects, respectively ^50^). Welch’s t-tests were conducted to assess sex differences in ADI-R and BRIEF scores. As here we were examining a series of tests and hypothesizing that these groups were not significantly different, we adjusted for 6 tests (critical p-value = 0.008); ^49,51^, with values less than 0.05 considered on the border of statistical significance. A chi-squared statistic was calculated to assess the frequency distribution of comorbid ADHD between sexes. For the t-tests, Glass’ delta—which is unaffected by unequal variances—was used as a measures of effect size.

To assess the association between ADI-R sub-scores (i.e., reciprocal social interaction, communication, and restricted, repetitive and stereotyped behavior) and EF (BRIEF GEC), we first calculated a Pearson correlation coefficient. To assess the impact of covariates (i.e., sex, IQ, age, ADHD, and a sex * EF interaction) on the association between ADI-R sub-scores and BRIEF GEC, we fitted a series of nested multiple regression models and then compared the fit of these models by calculating Akaike information criterion (AIC) values and F-ratios for model change. Lower AIC values are indicative of better model fit. As we were interested in three sub-scores from the ADI-R for these multiple regression models, we adjusted the critical value for 3 tests (critical p-value = 0.017), with values less than 0.05 considered on the border of statistical significance for the purposes of these analyses. Although this is an arbitrary cutoff for values considered to be on the border of statistical significance, we chose 0.05 as this is the value traditionally used when not corrected for multiple comparisons. To generalise the regression results beyond the given samples, robust regression was performed in the event of non-normally distributed standardized residuals via bootstrapping with 2000 samples. We obtained bootstrapped 95% confidence intervals for the model intercept and slopes and compared these with the confidence intervals from the original model. Similar confidence intervals between original and bootstrapped models would suggest that there are no considerable problems with non-normal distribution of residuals in the original models. Finally, we assessed the relationship between BRIEF GEC and ADI-R sub-scores in the male and female subgroups and Fisher’s *z* test was used to assess whether these correlations were significantly different. To examine the impact of more closely matched boys and girls on age and IQ, the same model fit and comparison procedure was performed on a subset of the sample, which was generated using the FUZZY extension command in SPSS. These analyses can be found in the supplement section. We allowed cases to be matched on age within 2 years and total IQ within 10 points. Three girls had missing full-scale IQ data, so the 22 girls with no missing values were matched to 44 boys.

## Results

### Sex differences in age, IQ, ADI-R scores, and BRIEF scores

There were no statistically significant differences between sexes (critical alpha adjusted to *p* = .008) in any of the ADI-R domains, BRIEF GEC, full-scale IQ, or age (Table 1). However, there were tendencies for girls to be slightly older (*p* = 0.029), have some more difficulties on the BRIEF index MI (*p* = 0.045) and to have less difficulties with the ADI-R C domain restricted and repetitive behaviour (*p* = 0.038) than the boys, but these sex differences did not reach the adjusted significance level.

**Table 1.**
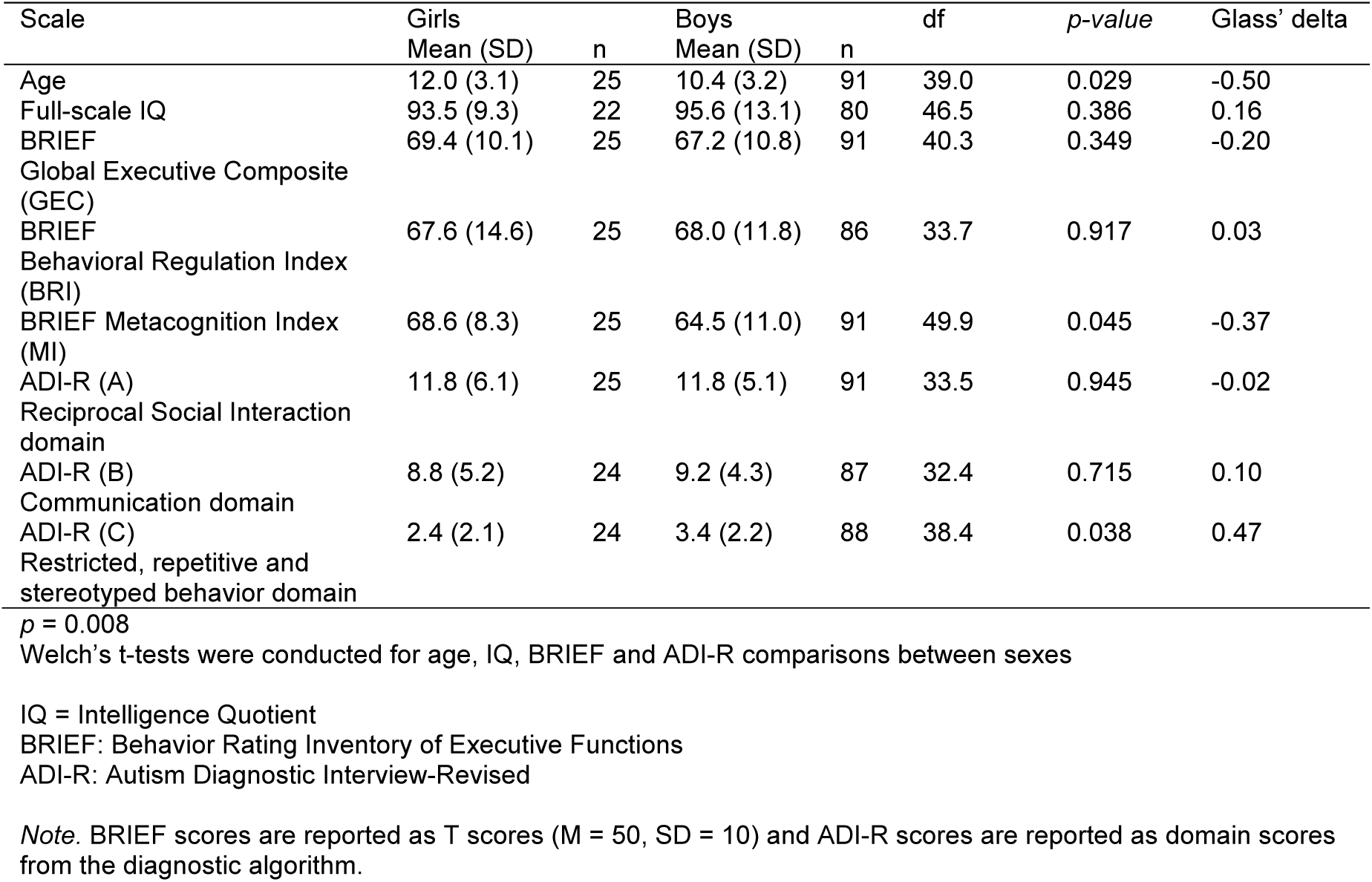
Age, IQ, BRIEF and ADI-R scores for girls and boys with ASD (N=116)

There was no significant diffference in the proportion of males and females with comorbid ADHD (χ^2^ = 2.96, *p* = 0.09).

### The association between reciprocal social interaction and executive function

There was a statistically significant correlation (adjusted critical alpha = 0.017) between reciprocal social interaction and EF (r = 0.31, *p* < 0.001), as indexed by scores on the ADI-R-A and BRIEF GEC, respectively. We fitted three nested linear regression models to assess the role of covariates (i.e., sex, IQ, age, and ADHD diagnosis) and the interaction of sex and EF on the relationship between reciprocal social interaction and BRIEF GEC (Table 2A). The first model, which included sex, IQ, age, and ADHD diagnosis, was not statistically significant (*p* = 0.49). The second nested model, which added BRIEF GEC, was on the border of our adjusted statistical significance threshold (*p* = 0.04). The second model (AIC = 630.9) was a significantly better fit of the data than the first model (AIC = 637.4; F(1, 96 = 8.38, *p* = 0.005), indicating that EF is related to reciprocal social interaction, over and above the main effects of sex, IQ, age, and ADHD diagnosis. The third nested model, which added the interaction of BRIEF GEC and sex, significantly predicted social interaction (*p* = 0.001). In this model, BRIEF GEC, sex, and their interaction provided a statistically significant contribution (Table 2A). The third model (AIC = 619.7), which included a sex * BRIEF GEC interaction term, was a significantly better model for the data than the second model, which only included main effects (AIC = 630.9; F(1, 95) = 13.15, *p* < 0.001).

**Table 2A-C.** Nested hierarchical models summary 2A Reciprocal Social Interaction domain

| ADI-R A | R<sup>2</sup> | B | SE B | p |
| --- | --- | --- | --- | --- |
| Model 1 | 0.03 |  |  | 0.486 |
| Constant |  | 19.98 | 5.20 | < .001* |
| Sex |  | -0.13 | 1.32 | 0.923 |
| IQ |  | -0.07 | 0.04 | 0.104 |
| ADHD diagnosis |  | -1.26 | 1.16 | 0.279 |
| Age |  | -0.08 | 0.17 | 0.618 |
| Model 2 | 0.11 |  |  | 0.041 |
| Constant |  | 10.68 | 5.95 | 0.076 |
| Sex |  | -0.55 | 1.28 | 0.667 |
| IQ |  | -0.07 | 0.04 | 0.106 |
| ADHD diagnosis |  | -1.88 | 1.14 | 0.102 |
| Age |  | -0.07 | 0.16 | 0.650 |
| BRIEF GEC |  | 0.14 | 0.05 | 0.005* |
| Model 3 | 0.22 |  |  | <.001* |
| Constant |  | 44.69 | 10.93 | <.001* |
| Sex |  | -29.20 | 7.99 | <.001* |
| IQ |  | -0.07 | 0.04 | 0.100 |
| ADHD diagnosis |  | -1.04 | 1.10 | 0.345 |
| Age |  | -0.05 | 0.15 | 0.743 |
| BRIEF GEC |  | -0.37 | 0.15 | 0.015* |
| BRIEF GEC * Sex |  | 0.42 | 0.11 | < .001* |

\* $p = 0.017$
ADI-R: Autism Diagnostic Interview- Revised, diagnostic algorithm. A: Reciprocal Social Interaction domain, B: Communication domain, C: Restricted, repetitive and stereotyped behavior domain.
ADHD: Attention deficit/ hyperactivity disorder
IQ: Intelligence Quotient
B = unstandardized regression coefficients

The standardized residuals from models 1 (*p* = 0.02), 2 (*p* = 0.01), and 3 (*p* = 0.003) were not normally distributed. Confidence intervals for the intercept and slopes of this model were similar to a bootstrapped model (Table S3A), indicating that there were no considerable problems with non-normal distribution of residuals in the model. The relationship between ADI-R A and BRIEF GEC was statistically significant in females (*p* < 0.001), but not males (*p* = 0.08; Figure 1). A formal comparison of these correlations suggested that the relationship between EF and reciprocal social interaction is stronger in females than males (Fisher’s z = − 3.56, *p* < 0.001). The same model fit and comparison procedure on subset of participants more closely matched on age and IQ revealed similar results (Supplementary material S2A).

**Figure 1.**
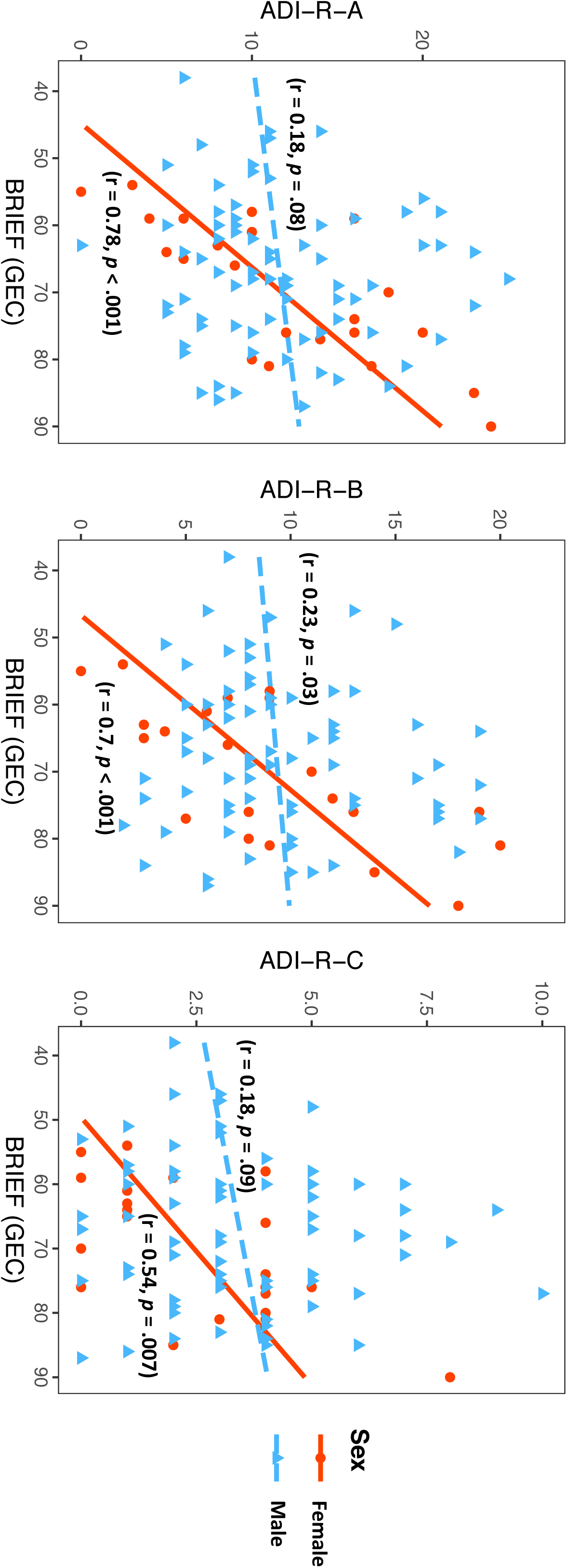
Correlations between ADI-R and BRIEF scores for girls and boys. ADI-R: Autism Diagnostic Interview-Revised, diagnostic algorithm. A: Reciprocal Social Interaction domain, B: Communication domain, C: Restricted, repetitive and stereotyped behavior domain. BRIEF_GEC: Behavior Rating Inventory of Executive Function, Global Executive Composite *Note:* BRIEF scores are reported as T scores (M = 50, SD = 10) and ADI-R scores are reported as domain scores from the diagnostic algorithm.

### The association between communication and executive function

There was a statistically significant correlation (adjusted critical alpha = 0.017) between communication and EF (r = 0.33, *p* < 0.001), as indexed by scores on the ADI-R-B and BRIEF GEC, respectively. We fitted three nested linear regression models to assess the role of covariates and the interaction of sex and EF on the relationship between ADI-R B and BRIEF GEC (Table 2B). The first model, which including sex, IQ, age, and ADHD diagnosis, was not statistically significant (*p* = 0.84). Although the second nested model was also not statistically significant (*p* = 0.20), BRIEF GEC provided a contribution that was on the border of statistical significance (*p* = 0.02). This second model (AIC = 577.3) was a better fit of the data than the first model (AIC = 581.5; F(1, 92) = 5.98, *p* = 0.02), indicating that EF is related to communication, over and above the main effects of sex, IQ, and ADHD diagnosis. However, this effect was on the border of statistical significance (*p* = 0.02) and needs to be validated in future studies. The third nested model, which added the interaction of BRIEF GEC and sex, significantly predicted communication (*p* = 0.004). In this model, BRIEF GEC, sex, and their interaction provided a statistically significant contribution (Table 2B). The third model (AIC = 566.9), which included a sex * BRIEF GEC interaction term, was a significantly better model for the data than the second model, which only included main effects (AIC = 577.3; F(1, 91) = 12.27, *p* = 0.001). The standardized residuals from models 1 (*p* = 0.004), 2 (*p* = 0.02), and 3 (*p* = 0.01) were not normally distributed. Confidence intervals for the intercept and slopes of this model were similar to a bootstrapped model (Table S3B), indicating that there were no considerable problems with non-normal distribution of residuals in the model. The relationship between BRIEF GEC and ADI-R B was statistically significant in females (*p* < 0.001), but not males (*p* = 0.03; Figure 1). A formal comparison of these correlations suggested that the relationship between EF and communication is stronger in females than males (Fisher’s z = −2.62, *p* = 0.01). The same model fit and comparison procedure on subset of participants more closely matched on age and IQ revealed similar results (Supplementary material S2B).

2B Communication domain

| ADI-R B | R<sup>2</sup> | B | SE B | p |
| --- | --- | --- | --- | --- |
| Model 1 | 0.01 |  |  | 0.843 |
| Constant |  | 12.86 | 4.47 | 0.005* |
| Sex |  | -0.42 | 1.14 | 0.717 |
| IQ |  | -0.04 | 0.04 | 0.357 |
| ADHD diagnosis |  | 0.44 | 1.01 | 0.664 |
| Age |  | 0.01 | 0.15 | 0.963 |
| Model 2 | 0.07 |  |  | 0.200 |
| Constant |  | 6.11 | 5.16 | 0.239 |
| Sex |  | -0.76 | 1.12 | 0.500 |
| IQ |  | -0.03 | 0.04 | 0.378 |
| ADHD diagnosis |  | -0.01 | 1.00 | 0.990 |
| Age |  | 0.01 | 0.14 | 0.949 |
| BRIEF GEC |  | 0.11 | 0.04 | 0.016* |
| Model 3 | 0.18 |  |  | 0.004* |
| Constant |  | 35.20 | 9.62 | < .001* |
| Sex |  | -25.36 | 7.10 | < .001* |
| IQ |  | -0.03 | 0.03 | 0.404 |
| ADHD diagnosis |  | 0.75 | 0.97 | 0.439 |
| Age |  | 0.01 | 0.13 | 0.916 |
| BRIEF GEC |  | -0.33 | 0.13 | 0.013* |
| BRIEF GEC* Sex |  | 0.36 | 0.10 | < .001* |

\*p = 0.017
ADI-R: Autism Diagnostic Interview- Revised, diagnostic algorithm. A: Reciprocal Social Interaction domain, B: Communication domain, C: Restricted, repetitive and stereotyped behavior domain.
ADHD: Attention deficit/ hyperactivity disorder
IQ: Intelligence Quotient
BRIEF\_GEC: Behavior Rating Inventory of Executive Function, Global Executive Composite
B = unstandardized regression coefficients

### The association between restricted, repetitive and stereotyped behavior and executive function

The correlation between restricted, repetitive and stereotyped behavior and EF, as indexed by scores on the ADI-R-C and BRIEF GEC respectively, was on the border of the adjusted critical alpha (r = 0.22, *p* = 0.019; adjusted critical alpha = 0.017). We fitted three nested linear regression models to assess the role of covariates and the interaction of sex and EF on the relationship between repetitive behavior and EF (Table 2C). The first model, which including sex, IQ, age, and ADHD diagnosis, was not statistically significant (*p* = 0.43). Nor was the second nested model which added BRIEF GEC (*p* = 0.12). This second model (AIC = 439.9)was a better fit of the data than the first model (AIC = 443.1; F(1, 93) = 5.08, *p* = 0. 03), but was on the border of statistical significance. The third nested model (adding the interaction of BRIEF GEC and sex) was not statistically significant (*p* = 0.06) (Table 2C). The third model (AIC = 438.4) was a better fit of the data than the second model (AIC = 439.9), but this was not statistically significant (F(1, 92) = 3.3, *p* = 0.07). The standardized residuals from models 2 and 3, which included the predictor of EF were normally distributed (*p* > 0.05), however, they were not normally distributed for the first model (*p* = 0.01). Confidence intervals for the intercept and slopes of this model were similar to a bootstrapped model (Table S3C), indicating that there were no considerable problems with non-normal distribution of residuals in the model. The relationship between EF and repetitive behavior was statistically significant in females (*p* = 0.007) but not statistically significance in males (*p* = 0.09; Fig 1). However, formal comparisons of these two correlations showed that they were not significantly different (Fisher’s z = −1.72, *p* = 0.09). The same model fit and comparison procedure on subset of participants more closely matched on age and IQ revealed similar results (Supplementary material S2C).

2C Restricted, repetitive and stereotyped behavior domain

| ADI-R C | R<sup>2</sup> | B | SE B | p |
| --- | --- | --- | --- | --- |
| Model 1 | 0.04 |  |  | 0.435 |
| Constant |  | 5.70 | 2.16 | 0.010* |
| Sex |  | -1.00 | 0.55 | 0.073 |
| IQ |  | -0.01 | 0.02 | 0.478 |
| ADHD diagnosis |  | -0.39 | 0.48 | 0.422 |
| Age |  | 0.01 | 0.07 | 0.889 |
| Model 2 | 0.09 |  |  | 0.118 |
| Constant |  | 2.69 | 2.50 | 0.286 |
| Sex |  | -1.15 | 0.54 | 0.037 |
| IQ |  | -0.01 | 0.02 | 0.506 |
| ADHD diagnosis |  | -0.58 | 0.48 | 0.229 |
| Age |  | 0.01 | 0.07 | 0.873 |
| BRIEF GEC |  | 0.05 | 0.02 | 0.027 |
| Model 3 | 0.12 |  |  | 0.063 |
| Constant |  | 1.33 | 4.88 | 0.037 |
| Sex |  | -7.62 | 3.60 | 0.037 |
| IQ |  | -0.01 | 0.02 | 0.533 |
| ADHD diagnosis |  | -0.39 | 0.49 | 0.428 |
| Age |  | 0.01 | 0.07 | 0.857 |
| BRIEF GEC |  | -0.07 | 0.07 | 0.309 |
| BRIEF GEC * Sex |  | 0.09 | 0.05 | 0.072 |

\* $p = 0.017$
ADI-R: Autism Diagnostic Interview- Revised, diagnostic algorithm. A: Reciprocal Social Interaction domain, B: Communication domain, C: Restricted, repetitive and stereotyped behavior domain.
ADHD: Attention deficit/ hyperactivity disorder
IQ: Intelligence Quotient
BRIEF\_GEC: Behavior Rating Inventory of Executive Function, Global Executive Composite
B = unstandardized regression coefficients

## Discussion

The main finding of the current study is that there are sex differences in the relationship between EF in everyday life and social difficulties related to ASD. We found a strong association between the BRIEF (GEC) scores and the ADI-R domains reciprocal social interaction and communication in girls, while these relationships were small and non-significant in boys. We did not find sex differences in the relationship between executive dysfunction and restricted and repetitive behaviors. These results have implications for understanding the different clinical manifestations of ASD in girls and boys. The findings indicate that girls and boys might have a different relationship between cognitive and behavioural phenotypes, which may provide novel information in search for different etiologies in girls and boys with ASD. Furthermore, it supports the notion that there may be different reasons for the behavioural problems related to ASD in girls and boys, with girls’ social and communicative challenges more strongly related to EF deficits. This could also help to develop sex-differentiated interventions.

Of particular note, we found evidence for a relationship between EF deficits and difficulties in the domains social reciprocity and communication, but not for the relationship between EF deficits and restrictive and repetitive behavior (RRB). This differs from previous studies, which found that EF difficulties were mainly related to RRB ^52,53^. However, these studies did not investigate the differences between girls and boys. On the other hand, Kenworthy and colleagues showed that EF deficits, measured with both performance tests and parental questionnaires, were related to all three components of the triad of impairment in ASD ^28^.

We did not find any statistically significant sex differences in the total amount of difficulties with social reciprocity or communication (ADI-R A and ADI-R B). However, we did observe that girls had slightly less reported problems related to RRB (ADI-R C), which is in line with previous studies ^54^. Results from the Simons Simplex Collection showed lower levels of restricted interests in girls ^55^, and others have found that girls with ASD have less RBB compared to boys, especially for high functioning girls ^56^.

The participants in our study did not significantly differ in the total amount of executive difficulties (GEC), but girls had higher scores (were slightly more impaired) than boys on the metacognitive index from the BRIEF. White and colleagues ^30^ reported that girls showed more EF difficulties in a matched sample of 78 girls and 158 boys with ASD and ADHD symptomatology. The BRIEF (GEC) scores for girls and boys from their study are similar to our results; however, in our study the difference in GEC scores between girls and boys did not reach the corrected level of significance. This might be due to a smaller sample size and a stricter control for multiple testing in our study.

We showed a strong link between EF deficits in everyday life and social dysfunction for girls with ASD. However, EF deficits seem to have a weaker association to social dysfunction for boys, which suggest that their social difficulties may have a different etiology. Despite not collecting any genetic information in our study, the finding is consistent with earlier studies suggesting that girls require a greater genetic load to manifest autistics symptoms, and that their cognitive and behavior characteristics tend to be more severe than boys when they are diagnosed ^57^. The main finding in our study is not that girls with ASD have more EF deficits than boys, but that the EF deficits are stronger linked to core ASD symptoms in girls. Our study only investigated the association between EF and social function, and does not give insight into the causal relationship between these two functional areas. Still, it is reasonable to argue that in girls, EF difficulties might drive social difficulties. This possible causal explanation should be further investigated in follow-up studies.

In typically developed children, girls appear to be more mature than boys, better at adapting to the classroom environment and more sociable ^58^. These differences may explain why girls tend to outperform boys in the early school years ^58^. Consequently, there tends to be different societal expectations of girls and boys in terms of social functioning. Girls with ASD might have more difficulties socially interacting with other girls, than boys with ASD have socially interacting with other boys ^59,60^. Thus, when EF is impaired in girls with ASD, it may have stronger negative effects on their social functioning because it requires more of their total cognitive resources.

Although the ADI-R together with the ADOS is considered to be the gold standard for assessing ASD ^61,62^, recent studies suggest that these diagnostic instruments may not be equally effective in identifying symptoms in both sexes. Beggiato and colleagues ^54^ investigated if the ADI-R items discriminate between males and females, and found that in two large cohorts the ADI-R was better at classifying males than females. They argue that because clinicians use diagnostic tools (like the ADOS and the ADI-R) that are not gender specific, it is likely that girls are underrepresented. Other screening instruments for autism symptoms like the Autism Spectrum Screening Questionnaire (ASSQ) and the Social Responsiveness Scale (SRS) have gender-specific items or different norms for boys and girls, to better to capture the “female phenotype” of autism ^54^. Thus, although girls and boys in our study have the same level of difficulties in social reciprocity and communication, they might have different expressions of autism symptoms in everyday life. We did not use the screening tools ASSQ or SRS because ADI-R is considered the gold standard measure of autism symptomatology. Further, ADI-R involves a clinical rating and not just parent reports, taking into account the clinical judgment.

In our study 34.5% of the children had a comorbid diagnosis of ADHD. Both ASD and ADHD are characterized by executive dysfunction, but the two disorders typically differ in terms of which subdomains of EF that are affected. Where individuals with ADHD usually have problems with inhibition, those with ASD are more likely to have difficulties with flexibility and planning ^63^. Recently, it is suggested that as many as 40-70% of children and adolescents with ASD have a comorbid diagnosis of ADHD ^19,64,65^. This complicates the picture regarding EF deficits, considering that the two disorders typically represent different aspects of EF deficits. In our study we did not have any significant sex differences in the distribution of ADHD. Furthermore, we included ADHD diagnosis as a predictor in our nested regression models (Table 2A-C). ADHD diagnosis did not have a significant contribution to the outcome measures related to social reciprocity, communication or RRB. We argue that it is important to include children and adolescents with comorbid ADHD in research on ASD, because ADHD is a common comorbid disorder in clinical populations. However, it is important to be aware of the possible influence ADHD might have on executive measures. Future research should combine the measurements used in this study with genetic information and/or neuropsychological testing to investigate sex differences in the relationship between EF and social difficulties in more depth.

### Potential clinical implications

The finding that executive dysfunction and social difficulties are highly related in girls but not in boys might be important for various aspects of clinical practice. Firstly, when girls present high scores on the ADI-R, it is reasonable to assess for executive difficulties and vice versa. Furthermore, because girls might have a higher risk for executive dysfunction in combination with their social difficulties, the finding can have implications for the choice of interventions. Following this argument, it is possible that girls (with the same amount of social difficulties as boys) will benefit more from EF interventions. Some existing programs that aim to enhance EF have shown to be effective on both social problems and EF ^66^. However, to our knowledge, research is yet to investigated whether this treatment may be more effective for girls than boys. Future studies need to consider that sex differences might influence the effect of interventions.

### Strengths and limitations of the study

The study consists of a clinically well-defined sample of children and adolescents with ASD. Even though we have a reasonable number of girls, the total number of girls is still relatively small. The participants were recruited from specialist health care services, which may limit the findings to the more severe conditions. Previous studies have shown that girls referred to specialist clinics have more severe problems than boys ^57^. The girls in our study were slightly older than the boys, but age was accounted for in the nested linear models. The BRIEF is based on parent’s own observations and evaluations of the child. This parental bias might have influenced the findings, but on the other hand, these instruments have been shown to be ecologically valid measurements of how the child functions in everyday life. We have used the *t*-score from the BRIEF in the analyses, which have age and gender “corrected” norms, since *t*-scores are commonly used in literature, as well as clinical practice, and it is important to understand how different clinical tools influences each other. Both the BRIEF and the ADI-R are based on information from parents and this might bias the findings. However, while the BRIEF is a questionnaire, the ADI-R is a clinical semi-structured interview, which involves a clinical rating. Together, they both give important information about a child’s behaviour.

Another reason for the sex difference in ASD prevalence might be that girls have a different phenotype. Currently, the established diagnostic practices and tools like the ADOS and the ADI-R are not constructed or adapted to measure the subtle difficulties that girls may present with, which differ from the typical presentation of ASD symptoms in boys. Lai and colleagues suggest this might be a circular phenomenon, since an ASD diagnosis is based on behavioral descriptions, and the most common diagnostic tools are largely validated on the classic male phenotype of autism behaviors ^1^.

## Conclusion

We report sex differences in the relationship between executive dysfunction and social difficulties in individuals with ASD. Our study found a strong relationship between difficulties with social reciprocity and communication and parent-rated executive dysfunction in girls, while the same relationship was not evident in boys. These results suggest potential underlying factors related to different manifestations of ASD in males and females, which may have clinical implications.

## Abbreviations

ADHD: Attention Deficit Hyperactivity Disorder
ADI-R: Autistic Diagnostic Interview-Revised
ADOS: Autism Diagnostic Observation Scale
ASD: Autism Spectrum Disorder
BRIEF: Behavior Rating Inventory of Executive Function
BRI: Behavioral Regulation Index
D-KEFS: Delis-Kaplan Executive Function System
EF: Executive Function
GEC: Global Executive Composite
IQ: Intelligence Quotient
MI: Metacognitive Index
PDD-NOS: Pervasive Developmental Disorder – Not Otherwise Specified
RRB: Restrictive and Repetitive Behavior

## Acknowledgements

We are thankful to all the BUPgen participants and partners. The study is part of the BUPgen Study group and the research network NeuroDevelop.

## Availability of data and materials

The datasets used and analyses in the current study are available from the corresponding author on reasonable request.

## Authors’ contributions

TT, TN, MGØ and OAA planned and designed the study. TT, MGØ, REH, AK, ALH and SH collected the clinical information. TT, TN and DQ analysed the data and interpreted the results. TT wrote the first draft of the manuscript. All the authors contributed to the manuscript and read and approved the final manuscript.

## Ethics approval and consent to participate

Written informed consent was obtained from a parent and/or legal guardian for all participants under the age of 18 years who were included in the study. Participants over 18 years gave written consent themselves. The study was approved by the Regional Ethical Committee and the Norwegian Data Inspectorate (REK #2012/1967), and was conducted in accordance with the Helsinki Declaration of the World Medical Association Assembly.

## Consent for publication

All the participants consented to publication.

