## Supplement Table 1-3 for "Sex differences in the relationship between social difficulties and executive dysfunction in children and adolescents with autism spectrum disorder"

Table S1. Age, IQ, BRIEF and ADI-R scores for girls and boys with ASD matched sample (N=66) [a]

| Scale | Girls | | Boys | | df | *p-value* | Glass' delta |
|---|---|---|---|---|---|---|---|
| | M (SD) | n | M (SD) | n | | | |
| Age | 11.9 (3.1) | 22 | 11.2 (3.3) | 44 | 44.6 | 0.415 | -0.21 |
| Full-scale IQ | 93.5 (9.3) | 22 | 92.1 (11.3) | 44 | 50.1 | 0.604 | -0.12 |
| BRIEF Global Executive Composite (GEC) | 69.5 (10.5) | 22 | 67.0 (11.3) | 44 | 45.0 | 0.376 | -0.22 |
| BRIEF Behavioral Regulation Index (BRI) | 68.3 (15.2) | 22 | 67.8 (11.7) | 42 | 34.6 | 0.891 | -0.04 |
| BRIEF Metacognition Index (MI) | 68.4 (8.7) | 22 | 64.2 (11.2) | 44 | 52.6 | 0.102 | -0.38 |
| ADI-R (A) Reciprocal Social Interaction domain | 11.7 (6.5) | 22 | 12.1 (5.1) | 44 | 34.4 | 0.842 | 0.08 |
| ADI-R (B) Communication domain | 8.9 (5.6) | 21 | 9.3 (4.5) | 43 | 33.2 | 0.803 | 0.09 |
| ADI-R (C) Restricted, repetitive and stereotyped behavior domain | 2.5 (2.1) | 21 | 3.9 (2.4) | 43 | 44.8 | 0.021 | 0.58 |

*p* = 0.008
Welch's t-tests were conducted for age, IQ, BRIEF and ADI-R comparisons between sexes.

IQ = Intelligence Quotient
BRIEF: Behavior Rating Inventory of Executive Functions
ADI-R: Autism Diagnostic Interview-Revised diagnostic algorithm
*Note.* BRIEF scores are reported as T scores (M = 50, SD = 10) and ADI-R scores are reported as domain scores from the diagnostic algorithm.

[a] The patients were recruited from Norwegian health services specializing in the assessment of ASD and other neurodevelopmental disorders. The study was part of the national BUPgen network. The matched sample consisted of 22 girls and 44 boys with ASD who were recruited between 2013 and May 2018 and assessed at age 5-19 years. Nine of the children (2 girls, 7 boys) were diagnosed with childhood autism, 5 (2 girls, 3 boys) with atypical autism, 27 (11 girls, 16 boys) with Asperger syndrome and 25 (7 girls, 18 boys) with unspecified pervasive developmental disorder (PDD-NOS). The male:female ratio was 2:1. In total, 17 children (25.8%) had a comorbid disorder of attention deficit/hyperactivity disorder (ADHD). Participants had an intelligence quotient (IQ) within the normal range based on a standardized Wechsler's test (Full-scale IQ ≥ 70) and spoke Norwegian fluently. Exclusion criteria were significant sensory losses (vision and/or hearing).

Table S2A-C Nested hierarchical models summary for the matched sample (N = 66)

S2A Reciprocal Social Interaction domain matched sample

| ADI-R A | | $R^2$ | *B* | *SE B* | *p* |
|---|---|---|---|---|---|
| Model 1 | | 0.03 | | | 0.740 |
| | Constant | | 20.95 | 6.80 | 0 .003* |
| | Sex | | -0.28 | 1.50 | 0.850 |
| | IQ | | -0.08 | 0.07 | 0.219 |
| | ADHD diagnosis | | -1.06 | 1.62 | 0.516 |
| | Age | | -0.06 | 0.22 | 0.783 |
| Model 2 | | 0.15 | | | 0.070 |
| | Constant | | 8.89 | 7.62 | 0.248 |
| | Sex | | -0.79 | 1.42 | 0.580 |
| | IQ | | -0.07 | 0.06 | 0.275 |
| | ADHD diagnosis | | -1.65 | 1.54 | 0.288 |
| | Age | | -0.10 | 0.21 | 0.614 |
| | BRIEF GEC | | 0.18 | 0.06 | 0.005* |
| Model 3 | | 0.29 | | | 0.002* |
| | Constant | | 46.82 | 13.16 | <0.001* |
| | Sex | | -30.27 | 8.75 | 0.001* |
| | IQ | | -0.07 | 0.06 | 0.203 |
| | ADHD diagnosis | | -0.43 | 1.46 | 0.768 |
| | Age | | -0.03 | 0.19 | 0.878 |
| | BRIEF GEC | | -0.39 | 0.18 | 0.031 |
| | BRIEF GEC * Sex | | 0,43 | 0.13 | 0.001* |

* $p = 0.017$
ADI-R: Autism Diagnostic Interview- Revised, diagnostic algorithm. A: Reciprocal Social Interaction domain, B: Communication domain, C: Restricted, repetitive and stereotyped behavior domain.
ADHD: Attention deficit/ hyperactivity disorder
IQ: Intelligence Quotient
BRIEF_GEC: Behavior Rating Inventory of Executive Function, Global Executive Composite
B = unstandardized regression coefficients

S2B Communication domain matched sample

| ADI-R B | | $R^2$ | *B* | *SE B* | *p* |
|---|---|---|---|---|---|
| Model 1 | | 0.04 | | | 0.604 |
| | Constant | | 16.87 | 5.89 | 0 .006* |
| | Sex | | -0.10 | 1.31 | 0.938 |
| | IQ | | -0.08 | 0.06 | 0.180 |
| | ADHD diagnosis | | 0.95 | 1.43 | 0.511 |
| | Age | | -0.05 | 0.19 | 0.802 |
| Model 2 | | 0.14 | | | 0.122 |
| | Constant | | 7.85 | 6.72 | 0.248 |
| | Sex | | -0.56 | 1.27 | 0.662 |
| | IQ | | -0.07 | 0.06 | 0.239 |
| | ADHD diagnosis | | 0.56 | 1.38 | 0.685 |
| | Age | | -0.09 | 0.19 | 0.614 |
| | BRIEF GEC | | 0.13 | 0.05 | 0.016* |
| Model 3 | | 0.27 | | | 0.004* |
| | Constant | | 40.29 | 11.68 | 0.001* |
| | Sex | | -26.08 | 7.86 | 0.002* |
| | IQ | | -0.07 | 0.05 | 0.195 |
| | ADHD diagnosis | | 1.66 | 1.32 | 0.212 |
| | Age | | -0.05 | 0.17 | 0.764 |
| | BRIEF GEC | | -0.35 | 0.16 | 0.029 |
| | BRIEF GEC * Sex | | 0.37 | 0.11 | 0.002* |

* $p = 0.017$
ADI-R: Autism Diagnostic Interview- Revised, diagnostic algorithm. A: Reciprocal Social Interaction domain, B: Communication domain, C: Restricted, repetitive and stereotyped behavior domain.
ADHD: Attention deficit/ hyperactivity disorder
IQ: Intelligence Quotient
BRIEF_GEC: Behavior Rating Inventory of Executive Function, Global Executive Composite
B = unstandardized regression coefficients

S2C Restricted, repetitive and stereotyped behavior domain, matched sample

| ADI-R C | | $R^2$ | *B* | *SE B* | *p* |
|---|---|---|---|---|---|
| Model 1 | | 0.08 | | | 0.285 |
| | Constant | | 6.40 | 2.86 | 0.029 |
| | Sex | | -1.38 | 0.64 | 0.034 |
| | IQ | | -0.01 | 0.03 | 0.690 |
| | ADHD diagnosis | | 0.02 | 0.69 | 0.972 |
| | Age | | 0.00 | 0.09 | 0.958 |
| Model 2 | | 0.16 | | | 0.067 |
| | Constant | | 2.26 | 3.29 | 0.494 |
| | Sex | | -1.59 | 0.62 | 0.013* |
| | IQ | | -0.01 | 0.03 | 0.835 |
| | ADHD diagnosis | | -0.15 | 0.67 | 0.823 |
| | Age | | -0.03 | 0.09 | 0.776 |
| | BRIEF GEC | | 0.06 | 0.03 | 0.023 |
| Model 3 | | 0.19 | | | 0.048 |
| | Constant | | 10.33 | 6.10 | 0.096 |
| | Sex | | -7.94 | 4.10 | 0.058 |
| | IQ | | -0.01 | 0.03 | 0.824 |
| | ADHD diagnosis | | 0.12 | 0.69 | 0.859 |
| | Age | | -0.02 | 0.09 | 0.864 |
| | BRIEF GEC | | -0.06 | 0.08 | 0.472 |
| | BRIEF GEC * Sex | | 0.09 | 0.06 | 0.123 |

* $p$ = 0.017
ADI-R: Autism Diagnostic Interview- Revised, diagnostic algorithm. A: Reciprocal Social Interaction domain, B: Communication domain, C: Restricted, repetitive and stereotyped behavior domain.
ADHD: Attention deficit/ hyperactivity disorder
IQ: Intelligence Quotient
BRIEF_GEC: Behavior Rating Inventory of Executive Function, Global Executive Composite
B = unstandardized regression coefficients

Table S3A Bootstrapped model ADI-R A

Model 1

| Predictors | Direct model | | Bootstrapped model | |
|---|---|---|---|---|
| | 2.50 % | 97.50 % | 2.50 % | 97.50 % |
| Intercept | 9.65 | 30.30 | 10.88 | 29.43 |
| Sex | -2.75 | 2.49 | -3.27 | 2.81 |
| IQ | -0.16 | 0.02 | -0.15 | 0.00 |
| ADHD | -3.56 | 1.04 | -3.59 | 1.14 |
| Age | -0.42 | 0.25 | -0.40 | 0.19 |

| Model 2 | Direct model | | Bootstrapped model | |
|---|---|---|---|---|
| Predictors | 2.50 % | 97.50 % | 2.50 % | 97.50 % |
| Intercept | -1.14 | 22.49 | -0.59 | 20.59 |
| Sex | -3.09 | 1.99 | -3.21 | 2.25 |
| IQ | -0.15 | 0.01 | -0.13 | 0.01 |
| ADHD | -4.13 | 0.38 | -4.30 | 0.60 |
| Age | -0.40 | 0.25 | -0.39 | 0.23 |
| BRIEF GEC | 0.04 | 0.24 | 0.05 | 0.25 |

Model 3

| Predictors | Direct model | | Bootstrapped model | |
|---|---|---|---|---|
| | 2.50 % | 97.50 % | 2.50 % | 97.50 % |
| Intercept | 22.99 | 66.39 | 26.46 | 62.74 |
| Sex | -45.07 | -13.34 | -42.59 | -14.55 |
| IQ | -0.15 | 0.01 | -0.14 | 0.00 |
| ADHD | -3.21 | 1.14 | -3.40 | 1.25 |
| Age | -0.36 | 0.25 | -0.34 | 0.25 |
| BRIEF GEC | -0.66 | -0.07 | -0.63 | -0.11 |
| BRIEF GEC * Sex | 0.19 | 0.65 | 0.21 | 0.61 |

ADI-R: Autism Diagnostic Interview- Revised, diagnostic algoritm. A: Reciprocal Social Interaction domain, B: Communication domain, C: Restricted, repetitive and steroetyped behavior domain
IQ: Intelligence Quotient
ADHD: Attention deficit/ hyperactivity disorder
BRIEF_GEC: Behavior Rating Inventory of Executive Function, Global Executive Composite

Table S3B Bootstrapped model ADI-R B

Model 1

| Predictors | Direct model | | Bootstrapped model | |
|---|---|---|---|---|
| | 2.50 % | 97.50 % | 2.50 % | 97.50 % |
| Intercept | 3.98 | 21.74 | 4.34 | 21.20 |
| Sex | -2.68 | 1.85 | -2.82 | 2.31 |
| IQ | -0.11 | 0.04 | -0.10 | 0.04 |
| ADHD | -1.57 | 2.45 | -1.64 | 2.53 |
| Age | -0.28 | 0.30 | -0.23 | 0.27 |

Model 2

| Predictors | Direct model | | Bootstrapped model | |
|---|---|---|---|---|
| | 2.50 % | 97.50 % | 2.50 % | 97.50 % |
| Intercept | -4.14 | 16.35 | -3.40 | 14.52 |
| Sex | -2.99 | 1.47 | -2.98 | 1.70 |
| IQ | -0.11 | 0.04 | -0.10 | 0.03 |
| ADHD | -2.01 | 1.98 | -2.16 | 2.17 |
| Age | -0.27 | 0.29 | -0.28 | 0.28 |
| BRIEF GEC | 0.02 | 0.19 | 0.02 | 0.19 |

Model 3

| Predictors | Direct model | | Bootstrapped model | |
|---|---|---|---|---|
| | 2.50 % | 97.50 % | 2.50 % | 97.50 % |
| Intercept | 16.08 | 54.31 | 19.23 | 52.03 |
| Sex | -39.47 | -11.26 | -38.80 | -12.33 |
| IQ | -0.10 | 0.04 | -0.09 | 0.03 |
| ADHD | -1.18 | 2.68 | -1.27 | 2.76 |
| Age | -0.25 | 0.28 | -0.24 | 0.25 |
| BRIEF GEC | -0.59 | -0.07 | -0.59 | -0.09 |
| BRIEF GEC * Sex | 0.15 | 0.56 | 0.17 | 0.55 |

ADI-R: Autism Diagnostic Interview- Revised, diagnostic algorithm. A: Reciprocal Social Interaction domain, B: Communication domain, C: Restricted, repetitive and stereotyped behavior domain
IQ: Intelligence Quotient
ADHD: Attention deficit/ hyperactivity disorder
BRIEF_GEC: Behavior Rating Inventory of Executive Function, Global Executive Composite

Table S3C Bootstrapped model ADI-R C

Model 1

| Predictors | Direct model | | Bootstrapped model | |
|---|---|---|---|---|
| | 2.50 % | 97.50 % | 2.50 % | 97.50 % |
| Intercept | 1.41 | 9.98 | 1.78 | 9.67 |
| Sex | -2.09 | 0.10 | -2.05 | 0.10 |
| IQ | -0.05 | 0.02 | -0.05 | 0.02 |
| ADHD | -1.35 | 0.57 | -1.25 | 0.68 |
| Age | -0.13 | 0.15 | -0.11 | 0.13 |

Model 2

| Predictors | Direct model | | Bootstrapped model | |
|---|---|---|---|---|
| | 2.50 % | 97.50 % | 2.50 % | 97.50 % |
| Intercept | -2.28 | 7.65 | -2.03 | 7.51 |
| Sex | -2.23 | -0.07 | -2.14 | -0.10 |
| IQ | -0.05 | 0.02 | -0.05 | 0.02 |
| ADHD | -1.54 | 0.37 | -1.47 | 0.49 |
| Age | -0.13 | 0.15 | -0.10 | 0.13 |
| BRIEF GEC | 0.01 | 0.09 | 0.00 | 0.09 |

Model 3

| Predictors | Direct model | | Bootstrapped model | |
|---|---|---|---|---|
| | 2.50 % | 97.50 % | 2.50 % | 97.50 % |
| Intercept | 0.64 | 20.01 | 1.92 | 19.60 |
| Sex | -14.77 | -0.47 | -14.28 | -1.56 |
| IQ | -0.05 | 0.02 | -0.05 | 0.03 |
| ADHD | -1.35 | 0.58 | -1.30 | 0.64 |
| Age | -0.12 | 0.15 | -0.09 | 0.14 |
| BRIEF GEC | -0.20 | 0.06 | -0.19 | 0.05 |
| BRIEF GEC * Sex | -0.01 | 0.20 | 0.00 | 0.19 |

ADI-R: Autism Diagnostic Interview- Revised, diagnostic algorithm. A: Reciprocal Social Interaction domain, B: Communication domain, C: Restricted, repetitive and stereotyped behavior domain
IQ: Intelligence Quotient
ADHD: Attention deficit/ hyperactivity disorder
BRIEF_GEC: Behavior Rating Inventory of Executive Function, Global Executive Composite
